# Ventral Hippocampal Infusions of C20:0 Ceramide Stimulate Microglia and Induce Anhedonia-like Behavior in Female and Male Rats

**DOI:** 10.1101/2022.04.22.489192

**Authors:** Lubriel Sambolín, Adariana Feliciano, Lizmarie Tirado, Cristina Suárez, Dariangelly Pacheco, Wilfred Fonseca, Anixa Hernández, Maria Colón, James Porter

## Abstract

Increased long-chain C20:0 ceramides have been found in the serum of patients with depression. Moreover, ceramides are linked with increased microglia reactivity and inflammatory cytokine production, which are associated with depression. Since ceramides can readily cross the blood brain barrier, peripheral C20:0 ceramides could enter the brain, activate microglia, and induce depressive-like behavior. In this study, we determined whether localized infusion of C20:0 ceramides into the ventral hippocampus (VH) of rats is sufficient to activate microglia and induce depressive-like behaviors. Adult male and female rats received infusions of C20:0 ceramides or vehicle solution every other day for 2 weeks. After the third infusion, C20:0-infused animals showed reduced sucrose preference suggesting anhedonia-like behavior. However, 4-6 days after the last infusion, animals infused with C20:0 ceramides did not show anhedonia-like or despair-like behavior suggesting the behavioral effect was reversible. After the final behavioral test, C20:0-infused animals had more Iba-1+ microglia in the VH. Despite the increased number of VH microglia, the expression of inflammatory genes was not increased. In conclusion, our data suggest that localized increases in C20:0 ceramides in the VH are sufficient to induce microglia activation and reversible anhedonia-like behavior.

## 1. INTRODUCTION

As central molecules in the metabolism of sphingolipids, ceramides contribute to cell membrane structure and signaling to trigger the cellular effects produced by cytokines (Ogretmen et al., 2004; Singh et al., 1998). High concentrations of long-chain ceramides (C16:0, C18:0, C20:0) have been found in patients with major depressive disorder (MDD) (Gracia-Garcia et al., 2011), and total ceramides are increased in animal models with depressive-like behavior (Gulbins et al., 2013; Oliveira et al., 2016). In addition, evidence suggests that microglia could play a role in depression (Sild et al., 2017). Microglia express cytokines and participate in synaptic transmission, synapse formation, and neuroplasticity which are all functions that are impaired in major psychiatric conditions such as MDD (Singhal et al., 1998; Wake et al., 2013; Panatier et al., 2006; Papouin et al., 2017). Furthermore, overactivation and increased amounts of microglia have been found in postmortem brain tissue from patients with disorders such as MDD and anxiety (Brites et al., 2015; Setiawan et al., 2015).

Little is known about the relationship between long-chain ceramides and microglial activation. The accumulation of ceramides can produce inflammatory or anti-inflammatory responses depending on the acyl chain length and can affect neuronal development and synaptic activity (Jung et al., 2013). Short-chain C8 ceramides inhibit the inflammatory reactions to lipopolysaccharide in microglia and induce the release of neurotrophic factors (Nakajima et al. 2002), suggesting that short-chain ceramides reduce the inflammatory phenotype of microglia (Jung et al. 2013). However, the effect of long-chain ceramides on microglia is unknown, but the effect is likely to be important since long-chain ceramides reduce brain slice viability (Monte et al., 2012).

In this study we examined if increased long-chain C20:0 ceramides in the VH are sufficient to induce the expression of microglia-related genes and depressive-like behaviors. To assess this, we infused long-chain C20:0 ceramides into the VH of adult male and female rats and evaluated anhedonia- and despair-like behavior. The male and female rats showed anhedonia-like behavior one day after the third C20:0 ceramide infusion. However, five days after the last C20:0 infusion, rats showed increased VH microglia but did not show evidence of anhedonia-like or despair-like behavior or increased expression of inflammation markers in the VH. Our findings suggest that C20:0 ceramides activate microglia and produce short-term anhedonia-like behavior without inducing long-term hippocampal inflammation or depressive-like behaviors.

## 2. MATERIALS AND METHODS

### 2.1 Animals

All animals used for behavioral experiments and tissue collection for molecular analysis were treated according to the legal and ethics requirements of the Institutional Animal Care and Use Committee (IACUC) from Ponce Health Sciences University (PHSU). Male and female 53 to 60-days-old Sprague Dawley rats were obtained from PHSU/PRI Animal Facilities and maintained in standard laboratory conditions (12 hr light/dark cycle, 25°C), with food and water provided *ad libithum*.

### 2.2 VH cannula implantation and long-chain C20:0 ceramide infusion

Adult female and male rats (post-natal day 53 - 60) were surgically implanted with bilateral cannulas in the VH (AP -5.6 mm; ML ± 4.5 mm; DV -7.5 mm), which were fixed in place using anchor screws and dental cement. After cannula implantation, rats recovered for two weeks, during which they received handling during the last seven days to habituate them to cannula manipulation and reduce stress during infusions. Fourteen days after the VH cannula implantation, male and female rats received seven infusions of either 100 µM long-chain C20:0 ceramides or a vehicle solution every other day (48 hours between each infusion). The C20:0 (Cat No. 10724, Cayman Chemical Company) ceramide solution was prepared by diluting the drug in alcohol to produce a 1 mM C20:0 ceramide stock solution in 100% ethanol. The stock solution was diluted in saline 0.9% to a final concentration of a 100 µM C20:0 ceramide and 10% ethanol prior to infusion. The vehicle solution was composed of 10% ethanol in saline 0.9%. Infusions were given at a rate of 0.5 µL per minute per side over a period of two minutes.

### 2.3 Behavioral Tests

#### 2.3.1 Sucrose preference test (SPT)

To assess anhedonia-like behavior, rats received a SPT 24 hours after every infusion of long-chain ceramides or vehicle solution for a total of seven sessions of the SPT. First, rats went through 4 days of acclimation to the sucrose solution. On days 12 and 13, male and female rats were exposed to two bottles of water in their home cage. On day 14, one of the bottles of water was switched to a bottle containing a 1% sucrose solution so that the animals acclimated to the sucrose. On day 15, the bottles were changed, so animals did not develop preference for the side where the bottles are located. The day of the test, rats were water and food deprived for 5 hours and then the test was performed. This test consisted of giving the rats access to a water bottle and to another bottle that contained 1% of sucrose solution for one hour. Sucrose solution consumption was measured for one hour and expressed as percent of sucrose intake over water intake to assess anhedonia-like behavior.

#### 2.3.2 Sucrose grooming test (SGT)

SGT also was used to assess anhedonia-like or motivational behavior of the rats. This test consisted of adding a 10% sucrose solution onto the rat’s snout and dorsal coat using an atomizer spray. The sucrose solution was sprayed 5 times along the dorsal coat of the animal and one time on the snout. After this, we measured the duration of grooming activity, which consists of animal grooming the face, extremities and dorsal and ventral areas of the body. This activity was analyzed during a five-minute interval from a video recording of the animal’s behavior. After each session, the cage was cleaned with alcohol and wiped dry for the next subject to be tested.

#### 2.3.3 Forced swim test (FST)

The FST assessed despair-like behavior in the rats by measuring the time spent immobile, swimming, or struggling. Immobility was considered when rats were practically immobile and only doing slight movements to keep the head above water surface. Swimming was evaluated when rats moved all their paws to stay afloat and struggling was considered when rats moved the anterior and posterior paws, but the movement of the upper paws break the water surface. In this test we put the rats in a cylinder (15.75” in height and 11.8” in diameter) filled with water to measure immobility, swimming and struggling time for 15 minutes to acclimate the animal to the apparatus and the environment. Twenty-four hours later, immediately before sacrifice, rats were subjected to the test during a five-minutes interval in which the whole session is recorded using the Any MAZE software. After finishing the forced swim test, rats were placed in a warm cage for complete drying and the cylinder water is changed between subjects.

### 2.4 Immunofluorescence

Iba-1 immunofluorescence staining was performed on paraffin-embedded tissue. After brain extraction, a sagittal cut was done and one hemisphere was fixed in 10% paraformaldehyde. Tissue was embedded in paraffin and coronal sections of VH were cut at 4□μm using a microtome. Slides were washed in xylene followed by hydration of tissue with ethanol (CDA-19). Slides were rinsed in 10% PBS and then heated while incubating with antigen retrieval 0.01□M Citrate-EDTA buffer (pH = 6.2) at 90–95□°C for 40□min. Tissues were washed with deionized water and remained in PBS for 5 minutes. Non-specific binding was prevented by adding protein block (Cat No. 50062Z, Life Technologies, Frederick, MD.) for 15□min. Tissues were incubated with an Anti-Iba-1 rabbit polyclonal primary antibody (Cat no. 019-19741, FUJIFILM Wako Pure Chemical Corporation) overnight in a humidified chamber at 4°C. A negative control was included by adding PBS instead of primary antibody. On the second day, slides were washed twice with PBS and incubated with Goat Anti-Rabbit IgG Secondary Antibody conjugated with Alexa Fluor 555 (Cat No. A21429, Invitrogen by Thermo Fisher Scientific) for 30□minutes at room temperature. This was followed by two PBS washes and incubation with DAPI for 5□minutes to label the nuclei. Tissues were washed with PBS twice, and slides were mounted with ProLong Gold antifade (Cat No. P36934, Invitrogen by Thermo Fisher Scientific). Two to three representative images per brain sample were taken using an Olympus System Microscope Model BX60 (Olympus Life Siences Solution). ImageJ software was used to measure the mean fluorescence intensity (Chompre et al., 2019) and to perform individual microglial cells counting. For the measurement of area fraction the values will be reported as mean ± SEM.

### 2.5 Quantitative PCR

Animals were sacrificed 20-30 minutes after the last behavioral task (5 minutes FST). After sacrifice the brain was extracted, one hemisphere was fixed in formaldehyde for histology assessment and the second hemisphere was used for qPCR analysis. For this, VH was dissected, and the tissue was homogenized by diluting with 700µL of TRIzol reagent and centrifuging 3 minutes using magnetic beads. RNA was isolated using miRNeasy Micro Kit from Qiagen (Cat No. 217084). An RNA concentration and purity was estimated using the Thermo Scientific NanoDrop 2000 to calculate appropriate concentration for cDNA synthesis as specified in the manufacture’s manual. Complementary DNA was synthesized using the iScript cDNA Synthesis Kit from Bio-Rad (Cat No. 1708891). Then, the cDNA samples were diluted by a 1:20 factor and RT-PCR was performed using iQ SYBR Green Supermix from BioRad (Cat No. 1708882) to evaluate mRNA expression of Iba-1 (microglial marker) and inflammatory genes (TNF-a, IKBA, TLR-4, NLRP3, HMBG-1). The RT-PCR sequences of each primer are shown in the **Supplementary Table S1**. These genes were normalized using GAPDH as a housekeeping gene.

## 3. STATISTICS

All data was reported as the mean ± standard error of the mean. Ordinary two-way ANOVA with Tukey’s multiple comparison test was used to analyze the sucrose preference test. Unpaired parametric t-test with Welch’s correction was used to compare between treatments, vehicle vs ceramide infused animals for the behavioral analysis, immunofluorescence staining quantification and mRNA expression analysis on VH. All analyses were performed using GraphPad Prism 9.2.0 for Windows.

## 4. RESULTS

### 4.1 Localized VH infusions of C20:0 ceramides induced short-term anhedonia-like behavior in male and female rats

To determine if the localized infusion of C20:0 ceramides into the VH of rats would cause the development of anhedonia-like behavior, adult male and female rats were implanted with cannulas in the VH and infused with 100 µM C20:0 ceramide or vehicle every other day for two weeks. Anhedonia-like behavior was assessed with the SPT 24 hours after each VH infusion of C20:0 ceramides as shown in the experimental timeline in **Figure 1A**. Animals infused with C20:0 ceramides showed less sucrose preference after the third infusion, suggesting an accumulative effect of C20:0 ceramides that can cause an increased anhedonia-like behavior (**Figure 1B**, F (6, 6) = 4.824, p=0.0385). Moreover, the animals infused with C20:0 ceramides showed a reduction in the average sucrose preference across the seven C20:0 infusions (**Figure 1C**, F (15, 13) = 4.037, p=0.0069) further suggesting that elevated C20:0 ceramides in the VH are sufficient to induce anhedonia-like behavior.

**Figure 1.**
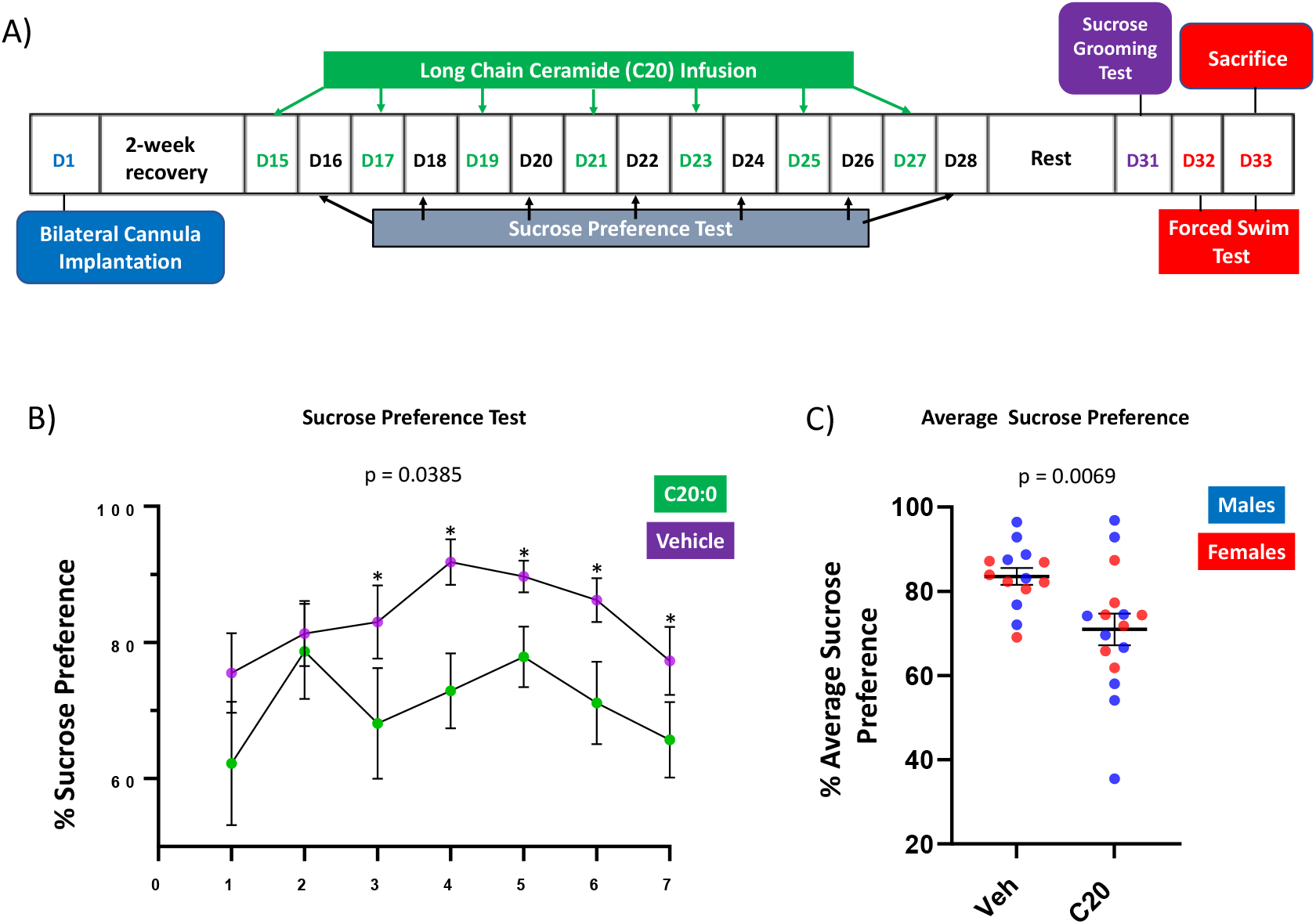
Long-chain C20:0 ceramide infusions into the VH induced anhedonia-like behavior in male and female rats. A) Timeline of the bilateral cannula implantation, C20:0 ceramide infusions every other day (green font), and intercalated sucrose preference tests 24 hours after every infusion (black font). D = day as indicated in the experimental timeline. B) Results of 7 sucrose preference tests given the day after each C20:0 ceramide infusion. C) Average sucrose preference across the 7 sucrose preference tests. n = 14 in vehicle group and n = 16 in ceramide group. Males in blue symbols and females in red symbols.

### 4.2 C20:0 ceramide infusions do not cause long-term anhedonia-like or despair-like behavior

Four to six days after the last assessment of sucrose preference, rats were exposed to the SGT to assess anhedonia-like and FST to assess despair-like behavior. The VH infusion of C20:0 ceramides did not affect the grooming activity (**Figure 2A**, F (9, 12) = 3.350, p=0.3685), suggesting that the anhedonia-like effect of the ceramides had reversed by four days after the last infusion. Like the SGT, the FST did not show differences in immobility (**Figure 2B**, F (10, 13) = 1.566, p=0.5662), swimming (**Figure 2C**, F (12, 9) = 3.275, p=0.4714), or struggling (**Figure 2D**, F (12, 9) = 1.303, p=0.3773). These results suggest that the localized infusion of C20:0 ceramides did not induce long-term depressive-like behaviors.

**Figure 2.**
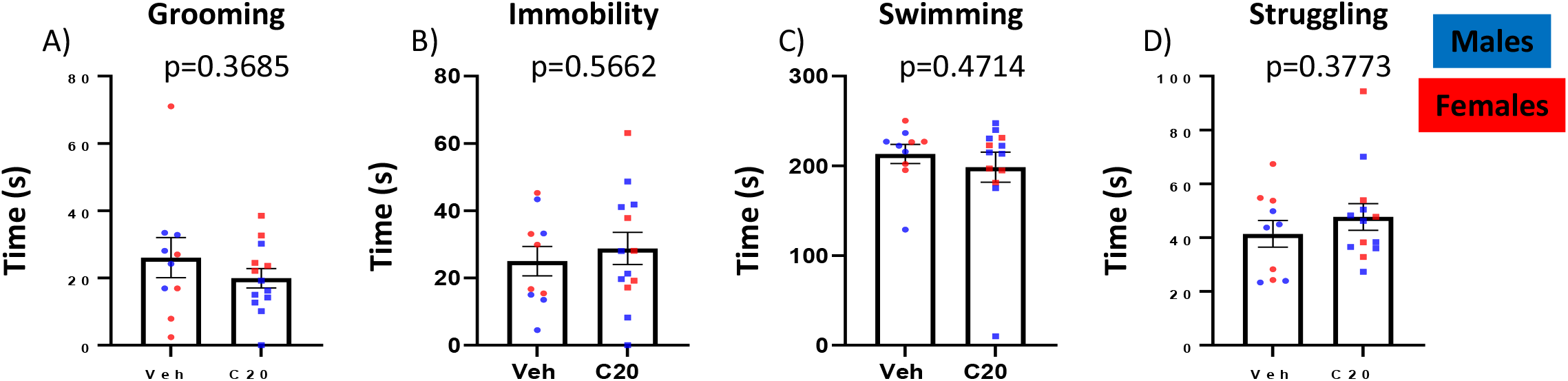
Long-term behavioral assessment four to six days after last C20:0 ceramide infusion. A) Sucrose grooming time during 5 minutes as an assessment of anhedonia-like behavior on day 31. B-D) Immobility, swimming, and struggling time measured in the forced swim test to assess despair-like behavior on day 33. N = 14 in vehicle group and n = 16 in ceramide group. Males in blue symbols and females in red symbols.

### 4.3 The infusion of C20:0 ceramides increased microglia in the VH

Since ceramides affect cultured microglia (Jung et al., 2013; Nakajima et al., 2002) and microglia are associated with anhedonia-like behavior (Henry et al., 2013), we investigated if the infusion of C20:0 ceramides affected microglia in the VH by labeling the microglial marker Iba-1 (**Figure 3A**). The infusion of C20:0 ceramides did not affect total Iba-1 protein expression measured as the percent of area labeled (**Figure 3B**, F (13, 15) = 1.318, p=0.6814). However, the rats that received the infusion of C20:0 ceramides showed more Iba-1+ microglia in the VH five days after the last infusion of C20:0 ceramides (**Figure 3C**, F (15, 13) = 1.930, p=0.0006) suggesting that the C20:0 ceramides increased the amount of microglia.

**Figure 3.**
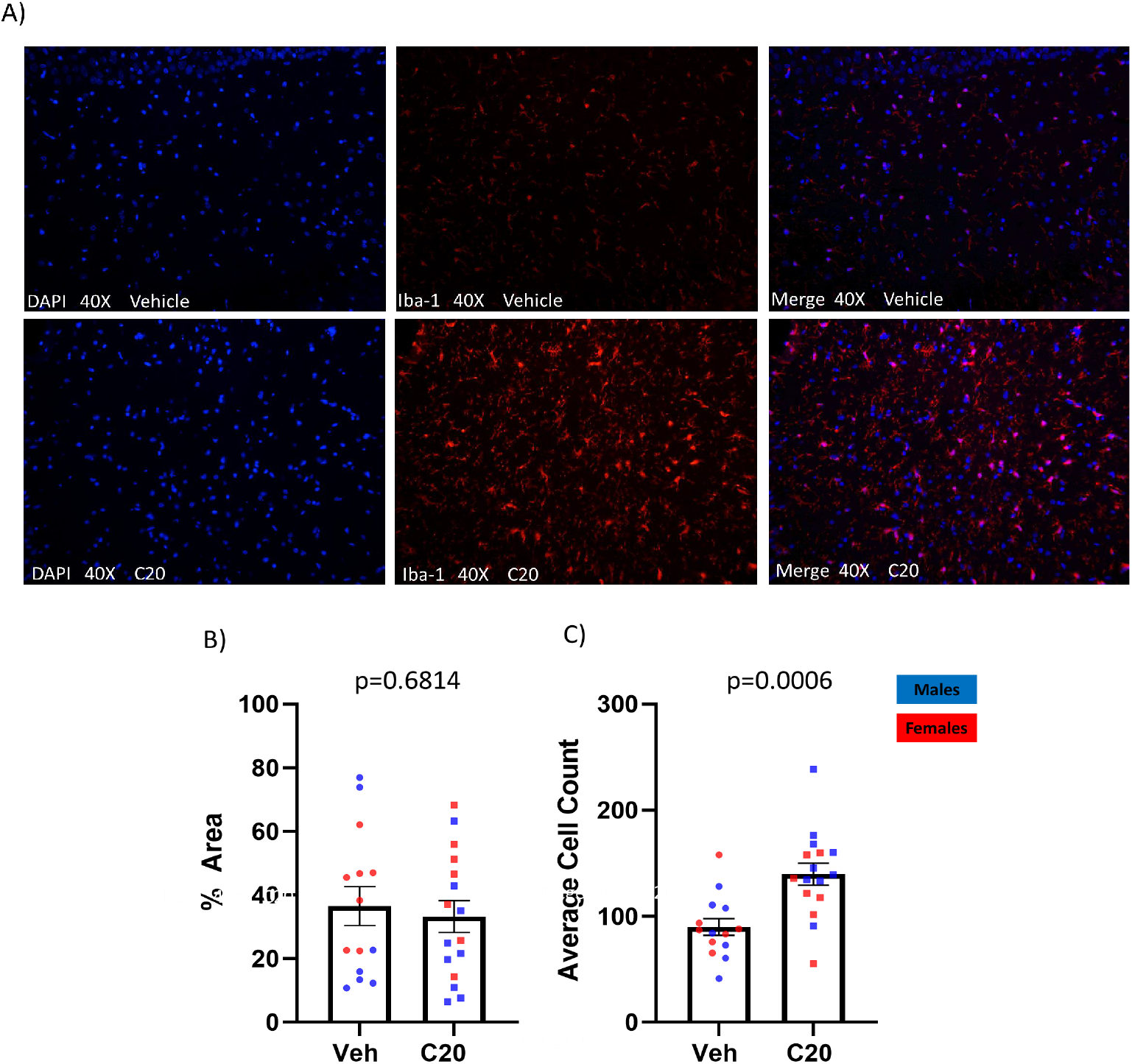
The VH contains more microglia six days after last C20:0 ceramide infusion. A) Representative images of VH from vehicle-infused (top row) and ceramide-infused (bottom row) rats showing cell nuclei labeled with DAPI and microglia labeled with Iba-1. B-C) The expression of the microglial marker Iba-1 in the VH of male and female rats quantified as percent of stained area and number of stained cells. N = 14 in vehicle group and n = 16 in ceramide group. Males in blue symbols and females in red symbols.

### 4.4 The infusion of C20:0 ceramides did not induce long-term neuroinflammation in the VH

In addition to quantifying the VH microglia, we also assessed whether the neuroinflammatory profile in the VH was increased five days after the last infusion of C20:0 ceramides. Consistent with the lack of change in overall Iba-1 protein expression, we found that the ceramide infusion did not change Iba-1 mRNA expression (**Figure 4A**, F (11, 9) = 2.282, p=0.1967). We found that the C20:0 infusions did not increase the mRNA expression of the inflammatory cytokine TNF-α (**Figure 4B**, F (9,11) = 8.909, p=0.7122) or the inflammasome NLRP3 (**Figure 4C**, F (11,9) = 1.852, p=0.2547). Moreover, we measured the mRNA expression of the high mobility group box-1 (HMGB-1), which can signal inflammation through activation of the inflammasome and NF-kB signaling cascade (Franklin et al., 2017; Crews et al., 2013; Scaffidi et al., 2002; Yang et al., 2009) and found difference in the ceramide-infused animals (**Figure 4D**, F (9, 11) = 18.40, p=0.2873). Lastly, we found that the infusion of ceramides did not affect the mRNA expression of the inflammatory modulator TLR-4 (**Figure 4E**, F (11,9) = 15.39, p=0.1060). Overall, the data suggests that the infusion of C20:0 ceramides into the VH did not induce long-term neuroinflammation.

**Figure 4.**
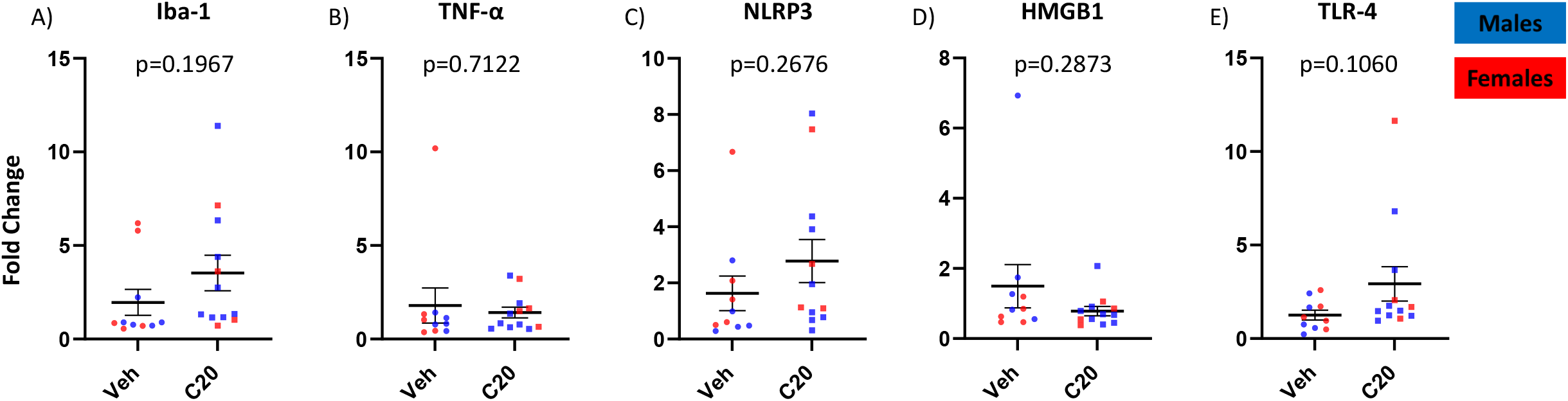
The mRNA expression of inflammatory genes in the VH six days after the last C20:0 ceramide infusion. A-E) Expression of Iba-1, TNF-α, NLRP3, HMGB1, and TLR-4 mRNA in VH tissue after infusion of C20:0 ceramides into the VH of male and female rats. N = 14 in vehicle group and n = 16 in ceramide group. Males in blue symbols and females in red symbols.

## 5. DISCUSSION

Although clinical findings suggest that long-chain C20:0 ceramides may contribute to depression (Gracia-Garcia et al., 2011; Hong et al., 2021; Brunkhorst-Kanaan et al., 2019), it is not clear whether increases in C20:0 ceramides in the brain are sufficient to induce depressive-like behavior in rodents. To address this gap in knowledge, we studied the effects of ventral hippocampal infusion of C20:0 ceramides on short- and long-term behavioral changes. We found that infusing C20:0 ceramides into the VH of female and male rats induced short-term anhedonia-like behavior beginning the day after the third infusion. However, the behavioral effects were transient. Four to six days after the last infusion, animals infused with C20:0 ceramide showed no signs of anhedonia-like or despair-like behavior or evidence of hippocampal inflammation. The C20:0 ceramide appears to have activated the microglia, since more microglia were found in the VH after C20:0 infusions. These findings suggest that increasing C20:0 ceramides in the VH induced transient anhedonia-like behavior.

Research suggests that long-chain ceramides in the hippocampus contribute to depressive-like behaviors in rodents. A previous study found strong evidence that fluoxetine and amitriptyline produce their antidepressant effects by reducing acid sphingomyelinase production of hippocampal ceramides in male mice (Gulbins et al., 2013). Furthermore, direct infusion of long-chain ceramides into the hippocampus are sufficient to induce depressive-like behaviors in rodents. For example, infusing C16:0 ceramides into the dorsal hippocampus caused anhedonia-like (Gulbins et al., 2013) and despair-like (Zoicas et al., 2019) behavior in male mice. We found that infusions of C20:0 ceramides into the VH also induced anhedonia-like behavior as determined by sucrose preference in both male and female rats. In contrast, infusion of C20:0 ceramides into either the VH in the current study or the dorsal hippocampus (Zoicas et al., 2019) did not affect despair-like behavior in the forced swim task. This suggests that the behavioral effects produced by increases in long-chain ceramides in the hippocampus may depend on the ceramide.

The increase in anhedonia-like behavior caused by the infused C20:0 ceramides was reversible. Rats showed more anhedonia-like behavior beginning the day after the third ceramide infusion. However, four to six days after the last infusion of C20:0 ceramides, ceramide-treated animals exhibited similar anhedonia-like behavior as the vehicle-treated animals. This suggests that the infusions did not produce lasting damage and the continued presence of the C20:0 ceramides is required to maintain the anhedonia-like behavior. In contrast, infusion of C16:0 ceramides into the dorsal hippocampus may cause longer-lasting behavioral effects, since reduced sucrose preference was observed up to four days after the last C16:0 infusion (Gulbins et al., 2013).

Although the anhedonia-like behavior reversed by four days, we still observed more VH microglia in the ceramide-treated animals suggesting that the C20:0 ceramides activated the microglia. In vitro and in vivo experiments show that short-chain ceramides reduced microglial reactivity and release of inflammatory cytokines (Nakajima et al., 2001, Nakajima et al., 2002, Jung et al., 2013). For example, pretreating isolated microglia with C2:0 ceramide reduced their inflammatory response to lipopolysaccharide (Jung et al., 2013). Since we did not find evidence of increased expression of inflammatory cytokines in the VH, the microglia did not seem to be producing inflammation but may be primed to produce larger inflammatory responses to subsequent stimulation with factors such as lipopolysaccharide (Jung et al., 2013; Scheiblich et al., 2017).

In conclusion, our findings suggest that the localized VH infusion of C20:0 ceramides was sufficient to induce short-term anhedonia-like behavior which reverted four days after the last infusion of ceramides. Furthermore, the C20:0 ceramides increased microglia in the VH but did not induce long-term neuroinflammation.

## Author contributions

**Lubriel Sambolin:** Conceptualization, Methodology, Investigation, Validation, Formal Analysis, Resources, Supervision, Writing – Original Draft, Writing – Review & Editing

**Adariana Feliciano:** Investigation, Formal Analysis, Writing – Original Draft,

**Lizmarie Tirado:** Investigation, Writing – Original Draft, Writing – Review & Editing

**Cristina Suarez:** Investigation, Writing – Review & Editing

**Dariangelly Pacheco:** Writing – Original Draft,

**Wilfred Fonseca:** Writing – Original Draft

**Anixa Hernandez :** Investigation

**Maria Colon :** Investigation

**James Porter:** Conceptualization, Methodology, Validation, Formal Analysis, Resources, Supervision, Writing – Original Draft, Writing – Review & Editing

## Acknowledgments

The authors acknowledge the involvement of graduate student Yesenia Rivera Escobales, Nashaly Irizarry Mendez, and Kimberly Santos Avilés. The authors would like to thank you Dr. Josue Pérez and Dr. Jaileene Pérez for the support with statistical analysis.

