## Supplemental table for "Ventral Hippocampal Infusions of C20:0 Ceramide Stimulate Microglia and Induce Anhedonia-like Behavior in Female and Male Rats"

| Gene | Name | Sequence |
|---|---|---|
| **GAPDH** | **Glyceraldehyde-3-Phosphate dehydrogenase** | F: 5'-GAT GCT GGT GCT GAG TAT GT-3'<br>R: 5'-GCG GAG ATG ATG ACC CTT T-3' |
| **IBA-1** | **Iodized calcium-binding adapter molecule 1** | F: 5'-CCT GAG GAG ATT TCA ACA GAA GC-3'<br>R: 5'-GGA CCG TTC TCA CAC TTC CC-3' |
| **TNF-α** | **Tumor necrosis factor alpha** | F: 5'-TCT TCT CAT TCC TGC TTG TGG C-3'<br>R: 5'-CAC TTG GTG GTT TGC TAC GAC G-3' |
| **HMGB-1** | **High-mobility group box 1** | F: 5'-GAG GTG GAA GAC CAT GTC TG-3'<br>R: 5'-AAG AAG AAG GCC GAA GGA GG-3' |
| **NLRP3** | **NLR family pyrin domain containing 3** | F: 5'-AGA AGC TGG GGT TGG TGA ATT-3'<br>R: 5'-GTT GTC TAA CTC CAG CAT CTG-3' |
| **TLR-4** | **Toll-like receptor 4** | F: 5'-TCC CTG CAT AGA GGT ACT TC-3'<br>R: 5'-CAC ACC TGG ATA AAT CCA GC-3' |

**Supplementary Table S1: Primer sequences for the assessment of mRNA levels of microglia cells and inflammatory markers.**
